# HAM-TBS: High accuracy methylation measurements via targeted bisulfite sequencing

**DOI:** 10.1101/163535

**Authors:** Simone Röh, Tobias Wiechmann, Susann Sauer, Maik Ködel, Elisabeth B. Binder, Nadine Provençal

## Abstract

**Background:** The ability to accurately and efficiently measure DNA methylation is vital to advance the understanding of this mechanism and its contribution to common diseases. Here, we present a highly accurate method to measure methylation using bisulfite sequencing (termed HAM-TBS). This novel method is able to assess DNA methylation in multiple samples with high accuracy in a cost-effective manner. We developed this assay for the *FKBP5* locus, an important gene in the regulation of the stress system and previously linked to stress-related disorders, but the method is applicable to any locus of interest.

**Results:** HAM-TBS enables multiplexing of up to 96 samples spanning a region of ~10 kb using the llumina MiSeq. It incorporates a triplicate bisulfite conversion step, pooled target enrichment via PCR, PCR-free library preparation and a minimum coverage of 1,000x. Furthermore, we designed and validated a targeted panel to specifically assess regulatory regions within the *FKBP5* locus including the transcription start site, topologically associated domain boundaries, intergenic and proximal enhancers as well as glucocorticoid receptor and CTCF binding sites that are not covered in commercially available DNA methylation arrays.

**Conclusions:** HAM-TBS represents a highly accurate, medium-throughput sequencing approach for robust detection of DNA methylation changes in specific target regions.

## Background

DNA methylation is the covalent addition of a methyl group at the 5-carbon ring of cytosine resulting in 5-methylcytosine (5mC). In the mammalian genome, this occurs predominantly in the context of CpG dinucleotides. It is one of several epigenetic marks influencing gene expression and serving multiple other purposes such as genomic imprinting, X chromosome inactivation and maintenance of genomic stability [1, 2]. Although 5mC is tightly regulated during cellular development, it has been shown to be responsive to external cues such as exposure to glucocorticoids and childhood maltreatment [3, 4, 5]. Aberrant regulation of the establishment, maintenance, erasure or recognition of DNA methylation has been associated with a range of disease phenotypes [6, 7]. The need to measure DNA methylation in large human cohorts in a cost-effective manner is therefore of paramount interest for research in epidemiology and psychiatry. Such advances would allow researchers to better assess the role of DNA methylation in relation to diseases as well as environmental impact on health. Indeed, there is increasing evidence in psychiatric disorders, that environmental risk factors and their interaction with genetic risk factors (GxE) are embedded via epigenetic changes, such as DNA methylation [3, 8]. Assessing DNA modifications with high accuracy and sensitivity in candidate loci would increase the power to detect and replicate such effects as well as perform time course experiments in large numbers of samples to understand the stability of the environmentally induced changes during development. In addition, changes related to environmental exposure such as adverse life events and psychopathology are often only present in specific cell types, although most studies rely on tissues such as post-mortem brain or blood samples. Assessing these effects in mixed tissues requires high accuracy in order to detect small changes emerging from a small number of cells. DNA bisulfite treatment followed by next generation sequencing enabled the quantification of DNA methylation marks at single base resolution. However, genome-wide bisulfite sequencing, although the best approach to identify DNA modifications, is still too cost intensive to be applied to large human cohorts at the coverage needed (˃60x) to detect differentially methylated sites [9]. Another set of accurate and cost-efficient measurement methods for DNA methylation at single CpG level are Illumina DNA methylation arrays. However, the ones currently available lack coverage in key enhancer regions that are important for environmentally-driven changes and have a relative small number of probes (~10-13) covering each site. Targetted bisulfite sequencing (TBS) offers a candidate approach to perform such studies with high resolution by increasing depth of read coverage per CpG to detect small changes in DNA methylation in a cost-efficient manner. Recently, few applications of TBS have been developed with differences in accuracy, throughput and library preparation [10, 11, 12, 13].

Our TBS approach focused on the *FKBP5* gene, which encodes the FK506 binding protein (FKBP51), a co-chaperon tightly involved in stress regulation. Genetic and epigenetic factors have repeatedly been shown to increase the activity of this gene and associated with increased stress-reactivity and psychiatric disorders [14]. We have previously reported allele-specific demethylation of CpG sites located in intronic enhancer regions of *FKBP5* specific to post-traumatic stress disorder (PTSD) in patients who had experienced child abuse [3]. These gene x environment interactions (GxE) may be mediated by differential susceptibility to adversity-induced changes in DNA methylation in specific enhancers. Current methods do not cover the relevant enhancer regions of *FKBP5* affected by stress exposure. A highly accurate, cost and time efficient method to investigate *FKBP5* DNA methylation in a large number of samples, is thus critical to gain more insight into how DNA methylation changes may mediate these GxE. In this manuscript, we present a cost-effective, high accuracy methylation measurement TBS (HAM-TBS) method to assess the regulatory regions of the *FKBP5* locus. Incorporating a triplicate bisulfite conversion step, PCR-free library preparation and rigorous quality control (validation of PCR target sites, *>*95% bisulfite conversion efficiency and 1,000x coverage minimum), ensures our method is extremely robust (Fig. 1). Medium throughput and handling accuracy of up to 96 samples spanning approximately 10 kb is facilitated by embedding the Hamilton pipetting robot and TapeStation with the Illumina MiSeq sequencer.

**Figure 1:**
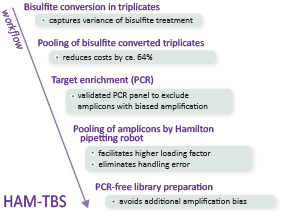
Workflow of the HAM-TBS method, depicting important processing steps and their advantages.

## Results

### QC, validation and optimization of the HAM-TBS method

TBS is based on bisulfite conversion coupled with targeted enrichment via PCR, library preparation for sequencing and subsequent quantification of methylation levels. All steps are necessary and may influence the outcome by introducing bias to the assessment of methylation levels or by insufficient quality control of the data. The standard approach to minimize potential biases before sequencing is to produce replicates and assess the mean methylation levels during the analysis. In order to design a highly accurate yet cost-effective and multiplexable approach, we needed to assess at which step (bisulfite conversion or amplification) and to what extent technical variability would be introduced, as well as which quality control steps to perform on the sequencing data to ensure a robust analysis. To this end, we assessed the methylation level of 0%, 25%, 50%, 75%, 100% *in vitro* methylated BAC control DNA for 3 different combinations of pooling strategies during the bisulfite treatment and PCR amplification (Fig. 2). Condition 1 (C1) assessed the methylation levels of control DNA using triplicate bisulfite treatments and PCR amplification for each replicate. C1 was considered the standard reference condition since each step was performed in triplicates. In condition 2 (C2) triplicate bisulfite treatments were pooled to perform one PCR amplification reducing the costs by approximately 64%. Finally, in condition 3 (C3) one bisulfite treatment of the control DNAs was performed followed by 3 separate PCR amplifications to assess the extend of the target enrichment bias. A smaller panel of 11 different PCRs (Fig. 3) within the *FKBP5* locus (Table S1) served as basis for this analysis.

**Figure 2:**
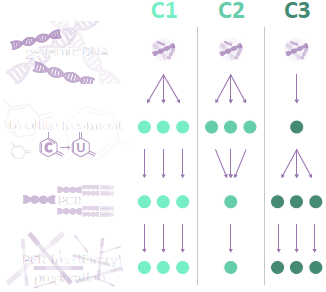
Setup of the TBS validation approach with the control conditions C1, C2 and C3. C1 is the reference condition with replicates in the bisulfite treatment and target enrichment step. C2 and C3 are more cost-effective versions dropping the replicate bisulfite treatment or target enrichment, respectively.

**Figure 3:**
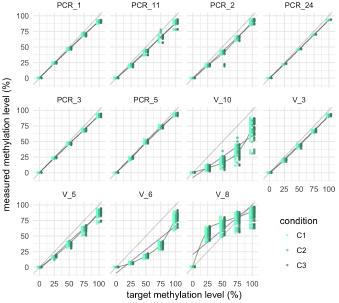
Methylation quantification of the control DNA used to evaluate the technical variability. Linear regression line (purple), loess fit line (green). PCR 3 was excluded due to low coverage, V 10 was excluded due to low coverage and non-linear amplification, V 6 and V 8 were excluded due to non-linear amplification.

Before comparing the three conditions, the collected sequencing data was subjected to three quality control steps in order to ensure accurate assessment of minimal methylation levels as well as small changes between samples.

1. **Bisulfite conversion rate** *>***95%.** We assessed the bisulfite conversion rate per sample and per amplicon and excluded rates lower than 95% from the analysis.
2. **Removal of PCR artefacts.** During the target amplification, the PCR occasionally introduces artefacts presenting non-existent CpG sequences in the target region. They present at very low coverage and extreme levels of methylation ~0 or ~100%). In order not to exclude potential SNPs giving rise to CpGs, we removed artefacts on this basis rather than limiting the analysis to known CpGs according to the reference genome.
3. **Minimum coverage of 1000x.** Higher sequencing depth and coverage of the CpGs yields higher accuracy of the methylation quantification. In order to determine the right balance between sequencing depth and thereby cost and sufficient accuracy, we took random samples of varying sequencing depth of a toy library and assessed the standard deviation for each level of methylation with respect to coverage (Fig. 4A). To find a meaningful cutoff for coverage, we considered the trade-off between sum of the average standard deviation per amplicon (cost) present in various levels of coverage (Fig. 4B). In concordance with [10], we identified 1000x coverage as a useful cutoff for our analysis, as the gain in accuracy with increasing coverage above this threshold is low and 1000x are reasonable to achieve for a larger locus, e.g. 9kb in the *FKBP5* panel.

All PCRs for our validation experiment showed bisulfite conversion levels *>*99%. After QC, a total of 40 CpG spread across 7 amplicons remained in our analysis (1 PCR failed due to coverage *<*1,000x, 1 showed non-linear amplification and coverage *<*1,000x, 2 showed non-linear amplification). Methylation levels were very similar between all 3 conditions with an average error of *<*1% when comparing absolute methylation levels of C2 and C3 versus C1 (Fig. 5B). We calculated the *R*^2^ values for each assessed CpG across the titration levels and used the mean per amplicon to compare the 3 conditions. *R*^2^ is a measure for assessing linearity of amplification of the methylation signal, which is crucial when quantifying methylation changes in e.g., cohort studies. Again, all conditions showed very high mean *R*^2^ values above 0.99 (Fig. 5A). This confirms that all conditions are suitable for high accuracy methylation detection. The introduced biases in our workflow based on the control DNA are minimal and enable very accurate methylation quantification even without including triplicates for the bisulfite conversion or target amplification. However, opposed to the target amplification, we cannot exclude slightly elevated variance of the bisulfite conversion on non-*in vitro* methylated DNA from e.g. patients. Therefore, we chose to use C2 for our HAM-TBS method. While it still maintains a triplicate bisulfite conversion step, it is the most cost-effective of the tested conditions, an important factor when processing many samples from cohort studies.

**Figure 4:**
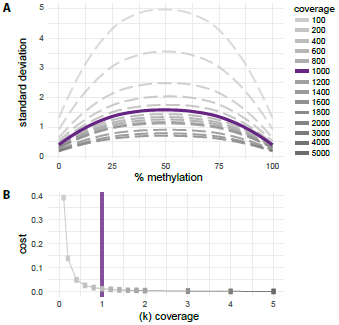
A:Standard deviation of varying coverage in respect to methylation level. **B:** Cost (accuracy as sum of the standard deviation) in respect to increasing coverage.

**Figure 5:**
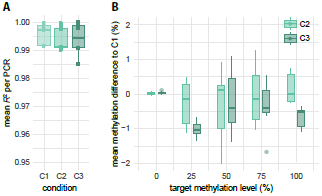
A:Mean *R*^2^ per amplicon for all conditions C1, C2 and C3. **B:** Mean methylation difference per PCR for C2 and C3 to the clean control condition C1. x-axis: target methylation levels during *in vitro* methylation.

### Comparison of the technical accuracy of pyrosequencing to TBS

Next, we aimed to compare TBS to pyrosequencing, the main method used for targeted DNA methylation analysis. Therefore, we assessed the methylation levels of 5 CpGs within PCR 5 and PCR 11 measured by pyrosequencing as well as in C1. The methylation analysis using pyrosequencing showed a high mean standard deviation of 4.68% with a maximum SD of 14.56%. The analysis using next generation sequencing with C1 showed a much lower mean standard deviation of 0.72% with a maximum SD of 1.83%. This demonstrates a significantly lower technical variation and therefore higher accuracy when assessing methylation levels using a TBS approach.

### Development of an extensive HAM-TBS *FKBP5* panel covering relevant regulatory sites

For future experiments investigating key regulatory sites of the *FKBP5* locus, we designed and validated a panel originally containing 32 amplicons. After QC, 29 of the amplicons were joint in the *FKBP5* panel comprising 315 CpGs (Table S2). The sequencing data did not show insufficient bisulfite conversion for any samples or 29 amplicons when performed on control DNA using C2. In total, 27 of the amplicons included in the panel presented good linearity (Sup. Figure 1) across the assessed methylation levels. Two amplicons showed very low methylation levels in the *in vitro* methylated control DNA (PCR 7, PCR 9). These amplicons are located near the transcription start site of *FKBP5* and have a very high CpG content. Therefore, we were not able to properly *in vitro* methylate the control DNA for this region and could not assess possible biases. To incorporate sites located in this region, these amplicons are included in the panel, but should be used with caution. PCR 26 of the HAM-TBS FKBP5 panel is located in the H19 locus [15] which is an imprinted gene and can serve as an internal positive control with a methylation level around 50% was performed.

## Discussion

We developed a targeted medium-throughput approach for measuring DNA methylation levels in multiple samples in parallel. This method enables cost-efficient high-resolution methylation measurements of target loci in cohorts of patients and probands at the *FKBP5* gene, a locus with large interest in the psychiatric and psychological community [14]. This cost-efficient, accurate method to determine *FKBP5* methylation levels would thus serve a large number of researchers. Our method is positioned between whole genome bisulfite sequencing and targeted approaches as pyrosequencing. The first is expensive and yields lower coverage and accuracy of single CpGs, the latter only allows us to assess very small regions at a time. HAM-TBS enables the analysis of a targeted but larger region (~10kb) at high resolution and low costs. DNA methylation studies in large cohorts, investigating the impact of environment or association with disease status in mixed tissues, however necessitate high accuracy at single site resolution. In fact, TBS was able to resolve methylation levels with a mean accuracy of 0.72%. A high level of accuracy was maintained in more cost-efficient approaches using only one PCR amplification round. By pooling triplicate bisulfite treatments prior to PCR amplification, we can account for variance introduced by the bisulfite treatment but also reduce costs and hands-on time during the target amplification.

The accuracy of the method benefits from a PCR-free library preparation and rigorous quality control (prior evaluation of linear PCR amplification of the target site, bisulfite conversion efficiency *>*95% and read coverage minimum of 1,000x). Nonetheless, a proper assessment of possible amplification biases due to the choice of amplicon location in the design step is crucial. Some loci show non-linear amplification curves, which renders them inappropriate for methylation quantification. Adjustment of primer design and PCR conditions may help solve this issue but for some loci, bias assessment could not be conducted. For instance in CpG islands with high CpG density, we found that *in vitro* methylation is not achievable using M.Sss1 methyltransferase. Therefore, bisulfite conversion and amplification bias could not be assessed in these regions. Differential methylation results from these sites should be interpreted with caution and perhaps require additional replication. Additionally, reaching 1,000x coverage is an important step to provide high resolution on methylation changes [10]. However, accurate quantification and pooling of many amplicons across multiple samples while reaching sufficient coverage of all regions has limitations. In theory, the MiSeq can handle a much higher loading factor (amplicons x samples) of almost 20,000 (not regarding uneven pooling of libraries, filtering of reads due to low quality or high amounts of PhiX), a maximum of 2,500-3,000 has proven to be feasible with minimal drop-out rates. Assuming multiplexing of 96 samples and 25 amplicons at an average length of 400 bp, a region of approximately 10kb can be comfortably covered with this approach. Notably, we streamlined the method to handle loading factors *>*2,000 by implementation of Aligent’s Tapestation and a pipetting robot for quantification and pooling of amplicons. Besides the throughput, this improves the robustness of the workflow. Our approach is designed to match the Illumina MiSeq due to its ability to run for 600 cylces resulting in 300bp long paired-end reads. This enables full-length coverage of amplicons up to a length of 600bp. While our approach can be applied to other loci, when working with a different sequencer, such as the HiSeq, it will be necessary to design shorter amplicons due to the limits of the sequencing chemistry. In the past years, few TBS methods have been developed tailored to their respective scope of research. Differences range from the species of interest, e.g. rat [10] or human [11, 12], to multiplexing capacity, library preparation method or size of the target loci. Thus far, e.g. Bernstein et al. [12] allows a panel of 48 indices, to our knowledge the approach by Chen et al. [11] allows for the theoretically highest multiplexing rate of 1536 samples due to custom made barcodes while in practice only using 478 to date. In this case, the high multiplexing capacity comes at the cost of an additional PCR step potentially introducing additional bias. Moreover, increasing the number of samples needs to be weighed against the size of the target region in order to ensure sufficient coverage. We identified 1000x as an optimal cutoff in terms of accuracy and cost in conjunction with Masser et al. [10]. In the study by Chen et al. [11], they use 100x as minimum cutoff. Based on our *in silico* analysis (Fig. 4A), this implies less accurate quantification of the methylation levels. Besides the number of samples the size of the region of interest is also an important factor. The method by Masser et al. [10] has been applied to 2 amplicons (233 and 320 bp), while Chen et al. enable the assessment of larger loci around 10 kb comparable to the HAM-TBS approach. Lastly, amplification based library preparation methods have been adapted by most TBS approaches. At this point, HAMTBS utilizes a PCR-free library preparation to avoid adding amplification biases.

Finally, using the optimized HAM-TBS workflow, we designed a panel comprising 29 amplicons to accurately assess methylation within the *FKBP5* locus using HAM-TBS. This panel covers ~9kb and targets important regulatory regions of the *FKBP5* gene including the transcription start site, intergenic and proximal enhancers and topologically associated domain (TAD) boundaries including CTCF sites. The HAM-TBS method and the *FKBP5* panel present valuable tools for epigenetic studies in which a highly accurate assessment of methylation levels is critical such as GxE studies in psychiatric research. It allows cost-efficient quantification of methylation in larger cohorts with optimized hands-on time due to automatization.

## Methods

### Generation of *in vitro* methylated control DNA

All primers designed for bisulfite PCR were first tested on *in vitro* methylated DNA to assess amplification efficiency and bias. For PCRs within the *FKBP5* gene, an *in vitro* methylated BAC (RP11-282I23, BACPAC) was used to generate control DNA. For PCRs outside the *FKBP5* locus (PCR 26, PCR 34, PCR 35), genomic DNA extracted from whole blood was amplified using the REPLI-g Mini Kit (QIAGEN GmbH, Hilden, Germany) to generate unmethylated DNA. 100% methylated DNA was achieved using *in vitro* methylation with M.SssI methyltransferase. After a first incubation (3 *μ*g DNA, 0.5 *μ*l SAM (32 mM), 1 *μ*l M.SssI (20 U/*μ*l, 40 *μ*l NEB buffer 2 [10x], diluted with ddH2O up to 400 *μ*l) of 4 h at 37C, 1 *μ*l of M.SssI (20 U/*μ*l) and 1 *μ*l of SAM (32 mM) was added, and a second 4 h incubation was performed. Subsequently, the reaction was purified using the nucleotide removal kit (QIAGEN GmbH, Hilden, Germany). *In vitro* methylation was repeated with the eluted DNA for a second time. 25%, 50% and 75% methylated control DNA was obtained by mixing 0% and 100% DNAs. *In vitro* methylation of control DNA was checked via pyrosequencing.

### Bisulfite treatment of DNA

We used the EZ DNA Methylation Kit (Zymo Research, Irvine, CA) in column and plate format depending on the amount of DNA and throughput needed. Between 200 to 500 ng were used as input DNA and processed according to the manufacturer’s instructions. DNA was eluted twice in 10 *μ*l elution buffer which recovered over 90% of the input DNA after bisulfite conversion when using the column format. In order to quantify bisulfite treated DNA, we use a spectrophotometer with RNA quantification settings.

### Target enrichment and amplicon pooling

The amplification of target locations from converted DNA was achieved using the TaKaRa EpiTaq HS Polymerase (Clontech, Mountain View, CA; final concentration: 0.025 U/l), bisulfite specific primers (final concentration of each primer: 0.4 M) and a touchdown cycling protocol with 49 cycles (for more details see Sup. Table 3 and section HAM-TBS *FKBP5* panel). The amplicons of all PCR reactions were quantified using the Agilent 2200 TapeStation (Agilent Technologies, Waldbronn, Germany) and equimolar pooled with the Hamilton pipetting robot. After speed-vacuum and resuspension in 50 *μ*l, a double-size selection was applied using Agencourt AM-Pure XP beads (Beckman Coulter GmbH, Krefeld, Germany) to remove excess of primers and genomic DNA.

### Pyrosequencing

Methylation analysis by pyrosequencing of 5 CpGs covered within PCR 5 (CpG 35607969, CpG 35608022) and PCR 11 (CpG 35690280, CpG 35690318, CpG 35690365) was performed in triplicates on BAC control DNA. Bisulfite conversion of *in vitro* methylated control DNA was applied as described above. Target enrichment by PCR was achieved with a biotinylated reverse primer but otherwise performed as described above. Pretreatment of PCR amplicons was facilitated with the PyroMark Q96 Vacuum Workstation (QIAGEN GmbH, Hilden, Germany). Sequencing of *FKBP5* CpGs was performed on a PyroMark Q96 ID system using PyroMark Gold Q96 reagents (QIAGEN GmbH, Hilden, Germany) and sequencing primers according to [3]: P4 S1 (TTTGGAGTAGTAGGTTAAA) GRE3 S1 MPI (GGGAATTATGAGGTTG). The PyroMark Q96 ID Software 2.5 (QIAGEN GmbH, Hilden, Germany) was used for data analyses.

### Library preparation and sequencing

For library generation, Illumina TruSeq DNA PCR-Free HT Library Prep Kit (Illumina, San Diego, CA) was used according to the manufacturer’s standard protocol. Qubit 1.0 (Thermo Fisher Scientific Inc., Schwerte, Germany) was used for quantification, Agilent’s 2100 Bioanalyzer (Agilent Technologies, Waldbronn, Germany) for quality assessment and Kapa HIFI Library quantification kit (Kapa Biosystems Inc., Wilmington, MA) for final quantification before pooling. Libraries were pooled equimolarly. Sequencing of the libraries was performed on an Illumina MiSeq using Reagent Kit v3 (Illumina, San Diego, CA; 600 cycles) in paired-end mode, with 30% PhiX added.

### Sequencing data processing

First, read quality was verified using FastQC [16]. Adapter sequences were trimmed using cutadapt v.1.9.1 [17]. For alignment to a restricted reference of hg19 based on the PCR locations, Bismark v.0.15.0 [18] was used. Due to the 600-cycle sequencing chemistry, PCRs shorter than 600 bp produce overlapping paired end reads. Using an in-house developed perl script, we trimmed low-quality overlapping ends. Quantification of methylation levels in CpG and CHH context was performed using the R package methylKit [19] with a minimum quality score of 20. The methylation calls were subjected to 3 quality control steps. First, we considered CHH levels for each sample and excluded samples if the conversion was less than 95% efficient. Second, we filtered PCR artefacts introduced during PCR amplification which present at low levels of coverage at 0 or 100% methylation. Lastly, according to our coverage cutoff, we excluded CpG sites supported by less than 1,000 reads. Subsequent analysis comparing methylation levels from the conditions C1, C2 and C3 as well as data from pyrosequencing was performed in R.

### Coverage considerations

When performing a sequencing experiment, one will usually sequence part of the generated library and quantify the methylation levels on this basis rather than sequence the whole library to see the true level within. Therefore, each sequencing experiment corresponds to drawing a random subset of a certain size (sequencing depth) of the whole library. Depending on the sequencing depth, this will yield a different level of accuracy of the methylation levels. We created a dataset simulating CpGs methylated at levels from 0 to 100% supported by 100.000 fragments each. Using a bootstrapping approach, we drew 100 random subsets of varying (sequencing) coverage (100, 200, 400,‥, 2000, 3000, 4000, 5000) for each level of methylation and the standard deviation (SD) was calculated. As a proxy for the increase in accuracy versus increase in sequencing depth (costs), the combined SD was divided by the sequencing depth.

### HAM-TBS

***FKBP5* panel**

We designed 29 primer pairs (Sup. Table 2) using BiSearch [20, 21] targeting the *FKBP5* locus. Initially, 32 PCRs were included but 3 PCRs were not selected for the panel due to QC failure. Positions of amplicons covering glucocorticoid response elements (GREs) were selected from [3] and the ChIP-seq track from the ENCODE project [22]. Amplicons covering CTCF binding sites were selected using HI-C peaks [23] and CTCF-ChIA-Pet interactions and CTCF ChIP-Seq information from the ENCODE project [22]. Only primers without CpGs in their sequence were chosen. The selected amplicons ranged from 200 to 450 bp in length.

## Acknowledgments

This study was funded by the BMBF grant Berlin-LCS (FKZ 01KR1301B) to EB and an ERC starting grant (GxE molmech, grant 281338) within the FP7 funding scheme of the EU to EB and fellowship from Canadian Institute of Health Research (CIHR) to NP. Layout of preprint publication modified from Matthias Legrand (CC BY-NC-SA 3.0).

## Author contributions

Experimental design by TW, SR, NP, EBB. Wet lab work by TW, SS, MK. Analysis by SR. Preparation of the manuscript by SR, TW, NP, EBB.

## Supplements

**Figure S1:**
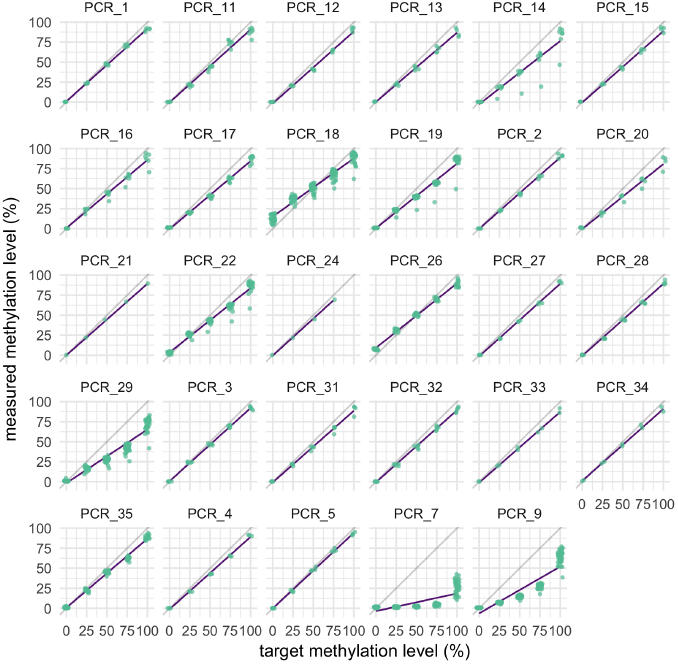
HAM-TBS *FKBP5* panel.

**Table S1:** Chromosomal location of validation panel. (#) indicates a primer published in [3]

| **PCR ID** | **chrom** | **start** | **stop** |
| --- | --- | --- | --- |
| V_3 (#) | chr6 | 35578699 | 35579119 |
| V_5 (#) | chr6 | 35655544 | 35655822 |
| V_6 (#) | chr6 | 35655792 | 35656028 |
| V_8 (#) | chr6 | 35656195 | 35656487 |
| V_10 (#) | chr6 | 35656760 | 35657069 |
| PCR_1 | chr6 | 35558361 | 35558652 |
| PCR_2 | chr6 | 35558459 | 35558774 |
| PCR_3 | chr6 | 35569680 | 35569946 |
| PCR_5 | chr6 | 35607754 | 35608065 |
| PCR_11 (#) | chr6 | 35690247 | 35690512 |
| PCR_24 | chr6 | 35570168 | 35570410 |
| PCR_26 | chr11 | 2023361 | 2023574 |

**Table S2:**
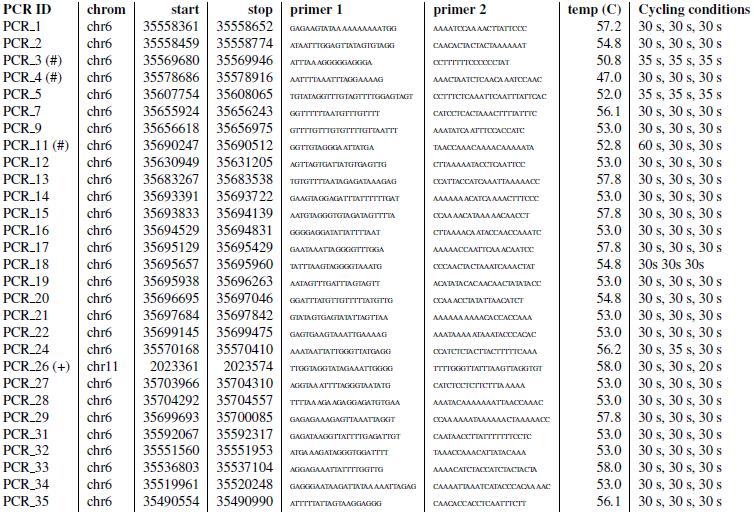
Location and primer sequences corresponding to HAM-TBS FKBP5 panel. (#) indicates a primer published in [3]; (+) indicates a primer published in [15].

## References

[1] A. Bird. Dna methylation patterns and epigenetic memory. Genes Dev, 16(1):6–21, 2002.

[2] M. Ehrlich and R.Y. Wang. 5-methylcytosine in eukaryotic dna. Science, 212(4501):1350–7, 1981.

[3] T. Klengel. Allele-specific fkbp5 dna demethylation mediates gene-childhood trauma interactions. Nat Neurosci, 16(1):33–41, 2013.

[4] R.S. Lee. A measure of glucocorticoid load provided by dna methylation of fkbp5 in mice. Psychopharmacology (Berl), 218(1):303–12, 2011.

[5] H. Thomassin. Glucocorticoid-induced dna demethylation and gene memory during development. EMBO J, 20(8):1974–83, 2001.

[6] A. Portela and M. Esteller. Epigenetic modifications and human disease. Nat Biotechnol, 28(10):1057–68, 2010.

[7] Y.L. Weng et al. Dna modifications and neurological disorders. Neurotherapeutics, 10(4):556–67, 2013.

[8] R. Yehuda et al. Holocaust exposure induced intergenerational effects on fkbp5 methylation. Biol Psychiatry, 80(5):372–80, 2016.

[9] M.J. Ziller et al. Coverage recommendations for methylation analysis by whole-genome bisulfite sequencing. Nat Methods, 12(230–2), 2015.

[10] D.R. Masser,A.S. Berg, and W.M. Freeman. Focused, high accuracy 5-methylcytosine quantitation with base resolution by benchtop next-generation sequencing. Epigenetics Chromatin, 6(1):33, 2013.

[11] D.R. Masser, D.R. Stanford, and W.M. Freeman. Targeted dna methylation analysis by next-generation sequencing. J Vis Exp., 24(96) 2015.

[12] D.L. Bernstein. The bispcr(2) method for targeted bisulfite sequencing. Epigenetics Chromatin, 8:27, 2015.

[13] G.G Chen et al. Medium throughput bisulfite sequencing for accurate detection of 5-methylcytosine and 5-hydroxymethylcytosine. BMC Genomics, 18(1):96, 2017.

[14] A.S. Zannas et al. Gene-stress-epigenetic regulation of fkbp5: Clinical and translational implications. Neuropsychopharmacology, 41(1):261– 74, 2016.

[15] C.C. Boissonnas et al. Specific epigenetic alterations of igf2-h19 locus in spermatozoa from infertile men. Eur J Hum Genet, 18(1):73–80, 2010.

[16] S.R. Andrews. Fastqc: A quality control tool for high throughput sequence data. Online Tool, 2010.

[17] M. Martin. Cutadapt removes adapter sequences from high-throughput sequencing reads. EMBnet journal, 17(1):10–12, 2011.

[18] F. Krueger and S.R. Andrews. Bismark: a flexible aligner and methylation caller for bisulfite-seq applications. Bioinformatics, 27(11):1571–2, 2011.

[19] A. Akalin et al. methylkit: a comprehensive r package for the analysis of genome-wide dna methylation profiles. Genome Biol, 13(10):R87 2012.

[20] T. Aranyi et al. The bisearch web server. BMC Bioinformatics, 7:431, 2006.

[21] G.E. Tusnady et al. Bisearch: primer-design and search tool for pcr on bisulfite-treated genomes. Nucleic Acids Res, 33(1):e9, 2005.

[22] E.P. Consortium. An integrated encyclopedia of dna elements in the human genome. Nature, 489(7414):57–74, 2012.

[23] S.S. Rao et al. A 3d map of the human genome at kilobase resolution reveals principles of chromatin looping. Cell, 159(7):1665–80, 2014.

